# The cerebellum acts as the analog to the medial temporal lobe for sensorimotor memory

**DOI:** 10.1101/2023.08.11.553008

**Authors:** Alkis M. Hadjiosif, Tricia L. Gibo, Maurice A. Smith

## Abstract

The cerebellum is critical for sensorimotor learning. The specific contribution that it makes, however, remains unclear. Inspired by the classic finding that, for declarative memories, medial temporal lobe structures provide a gateway to the formation of long-term memory but are not required for short-term memory, we hypothesized that, for sensorimotor memories, the cerebellum may play an analogous role. Here we studied the sensorimotor learning of individuals with severe ataxia from cerebellar degeneration. We dissected the memories they formed during sensorimotor learning into a short-term temporally-volatile component, that decays rapidly with a time constant of just 15-20sec and thus cannot lead to long-term retention, and a longer-term temporally-persistent component that is stable for 60 sec or more and leads to long-term retention. Remarkably, we find that these individuals display dramatically reduced levels of temporally-persistent sensorimotor memory, despite spared and even elevated levels of temporally-volatile sensorimotor memory. In particular, we find both impairment that systematically increases with memory window duration over shorter memory windows (<12 sec) and near-complete impairment of memory maintenance over longer memory windows (>25 sec). This dissociation uncovers a new role for the cerebellum as a gateway for the formation of long-term but not short-term sensorimotor memories, mirroring the role of the medial temporal lobe for declarative memories. It thus reveals the existence of distinct neural substrates for short-term and long-term sensorimotor memory, and it explains both newly-identified trial-to-trial differences and long-standing study-to-study differences in the effects of cerebellar damage on sensorimotor learning ability.

**Significance Statement:** A key discovery about the neural underpinnings of memory, made more than half a century ago, is that long-term, but not short-term, memory formation depends on neural structures in the brain’s medial temporal lobe (MTL). However, this dichotomy holds only for declarative memories – memories for explicit facts such as names and dates – as long-term procedural memories – memories for implicit knowledge such as sensorimotor skills – are largely unaffected even with substantial MTL damage. Here we demonstrate that the formation of long-term, but not short-term, sensorimotor memory depends on a neural structure known as the cerebellum, and we show that this finding explains the variability previously reported in the extent to which cerebellar damage affects sensorimotor learning.

## Introduction

The medial temporal lobe (MTL) – which includes the hippocampus – has been shown to serve as the gateway to long-term stable *declarative* memories, as MTL lesions severely impair long-term memory but largely spare short-term working memory^1–4^. For example, patient HM developed a severe deficit in long-term declarative memory formation following bilateral MTL resection, where he would be unable to remember that conversations even took place after they were over^1^. Yet, he could readily carry on a conversation while it lasted, showed normal performance in immediate recall tasks^3,5^, and could retain newly-acquired declarative information for as long as he could maintain continuous mental rehearsal^5^. Subsequent evidence has suggested that the dichotomy between the MTL’s role in long-term vs. short-term memory may not be absolute, as the MTL may play a role in some short-term declarative memories, with several studies reporting short-term declarative memory impairment in individuals with MTL damage^6–8^, and neurophysiologic studies showing that the MTL is active during short-term memory tasks^9–12^. Yet it remains clear that MTL damage has a greater effect on long-term than short-term declarative memory^7,13,14^. In contrast to the effects on the formation of declarative memories, however, MTL lesions can leave procedural learning largely intact, even over long timescales. Individuals with MTL lesions can readily acquire procedural memories for motor skills such as mirror tracing^15,16^, rotary pursuit^17^ and force-field (FF) adaptation^18^, and recall these memories after days or weeks^17^, or even a year^16^. Thus, individuals with MTL lesions may acquire a new skill with extensive practice, yet have no recollection of ever having practiced the skill^15^. If, in line with these findings, the MTL acts as a specific gateway to long-term declarative memory, then are there analogous structures that act as gateways to long-term procedural memory?

The cerebellum is critical for normal motor function^19–27^ and, more specifically, for the formation of sensorimotor memories^28–30^. In particular, it has been hypothesized that the cerebellum provides a substrate for adaptable internal models of the environment^31–36^. Correspondingly, cerebellar lesions in monkeys can eliminate the ability of eye saccades to adapt to weakened ocular muscles^26^ or to target shifts imposed during saccades^37^. Studies of cerebellar degeneration in humans have demonstrated deficits, not only in saccade adaptation^38,39^ but also across several other motor tasks, ranging from upper-limb visuomotor learning^40–47^ and FF learning^42,48–52^ to locomotor adaptation^53^ and speech adaptation^47,54^. There is, thus, ample evidence that the cerebellum is crucial for the formation of sensorimotor memory; however, we do not yet know whether it is differentially involved in the formation of long-term vs short-term sensorimotor memories.

A clue, perhaps, may lie in the differences between the severity of impairment reported in different studies of sensorimotor learning. For example, the level of impairment for sensorimotor learning varied considerably across three studies based on the same upper-limb FF adaptation paradigm. Interestingly, these studies employed very different trial spacing^49–51^, which would alter the duration for which memory needs to be maintained from one action to the next. More specifically, two of these studies used short, 5-7sec trial spacing and found only modest 30-40% deficits in sensorimotor learning^49,50^, whereas the third study used considerably longer, 25sec trial spacing and found a more profound 75% deficit^51^. This could suggest that tasks which require only short-term (5-7sec) maintenance of sensorimotor memories may mask the severity of cerebellar learning deficits that are more readily apparent when longer periods of maintenance are required. However, there were differences in other details of these experimental paradigms and in the patients who participated in each study, making it impossible to conclude that memory maintenance is the critical factor behind the observed differences in sensorimotor learning.

Here we overcome the limitations inherent in across-study comparison by taking advantage of the fact that, although the studies that showed only modest learning deficits in severe cerebellar ataxia had inter-trial intervals (ITIs) of only 5-7s on average, these intervals were not fixed for all trials but varied from one to the next. We, therefore, analyze the effects of these trial-to-trial variations in ITI on trial-to-trial differences in motor memory availability within each study. We hypothesize that cerebellar damage dramatically impairs the formation of longer-term temporally-persistent sensorimotor memory yet spares short-term temporally-volatile memory. If this were the case, it would lead to decreased temporally-persistent relative to temporally-volatile memory in individuals with severe cerebellar damage, resulting in an outsized reliance on short-term temporally-volatile memory that decays with a time constant of just 15-20s^55–57^. Our hypothesis would thus predict that (i) over short ITIs, impairment would rapidly increase with ITI, even for different trials from the same individual, and (ii) with large ITIs, a dramatic impairment would occur, comparable to the 75% deficit observed in the Smith and Shadmehr, 2005^51^ study. Remarkably, we find clear evidence for both predictions. As an additional test of this hypothesis, we then examined the findings from a larger group of 15 previous studies of sensorimotor learning in target-directed movements in individuals with cerebellar degeneration and found that those that employed a larger number of movement directions – increasing the effective ITI – reported dramatically increased impairment compared to those employing fewer movement directions. Together, these findings indicate that cerebellar degeneration impairs long-term sensorimotor memory but spares short-term memory, in line with the idea that the cerebellum may provide a gateway to the formation of long-term procedural memories, analogous to the role of the MTL for declarative memories.

## Results

Here we investigate whether the cerebellum acts as a gateway to long-term sensorimotor memory formation by examining the effects of cerebellar damage on the persistence of sensorimotor memory over the course of just 1-3 minutes or less, to determine whether short-term temporally-volatile and longer-term temporally-persistent memories are differentially impaired. To investigate this, we reanalyze the raw data from two different studies of FF adaptation during reaching arm movements that found only moderately impaired sensorimotor learning in individuals with severe cerebellar degeneration^49,50^, with the premise that the moderate impairment that was observed resulted from a heterogenous combination of unimpaired short-term memory and severely impaired longer-term memory. Specifically, we use the time intervals between movements, which were not originally examined, to dissect out temporally-volatile vs temporally-persistent memory based on the timespan of motor memory available on each trial. We then compare the ability to form each of these memories in healthy controls and patients with cerebellar disease.

We focused our analysis on patients with severe cerebellar disease (Severe-CBL group, n=20; 8 in the 2010 study and 12 in the 2013 study)), based on a threshold of 40 for the International Cooperative Ataxia Rating Scale (ICARS), matching that used in the Criscimagna-Hemminger et al., 2010 study. The Severe-CBL group was compared against age-matched controls (n=28; 13 in the 2010 study and 15 in the 2013 study). Note that, Criscimagna-Hemminger et al., 2010 separately analyzed individuals with milder (ICARS<40, n=5) and severe (ICARS≥40) cerebellar ataxia and found significant impairment only in the severe subgroup. We thus focus on patients with severe cerebellar ataxia throughout this paper; however, we also present analysis of patients with milder cerebellar ataxia in the Supplementary Materials (Figures S1, S2).

The task (see Methods) had participants make a series of center-out reaching arm movements, in which they were instructed to rapidly “shoot” through and beyond a target, positioned at one^50^ or two^49^ possible target locations while using a robotic manipulandum (Figure 1a). After a baseline period during which movements were not physically perturbed, participants received training where they were perturbed by a robotically-generated velocity-dependent lateral force-field (FF). This training lasted for 240 (2010 study) or 170 (2013 study) trials, during which rest breaks were present (three in the 2010 study and one in the 2013 study). Each individual was studied for two sessions, one where the FF was introduced abruptly and one where it was introduced gradually, counterbalanced in order, with the direction of the FF perturbation (clockwise vs counterclockwise) also counterbalanced in the Gibo et al., 2013 dataset but not in Criscimagna-Hemminger et al., 2010.

**Figure 1:**
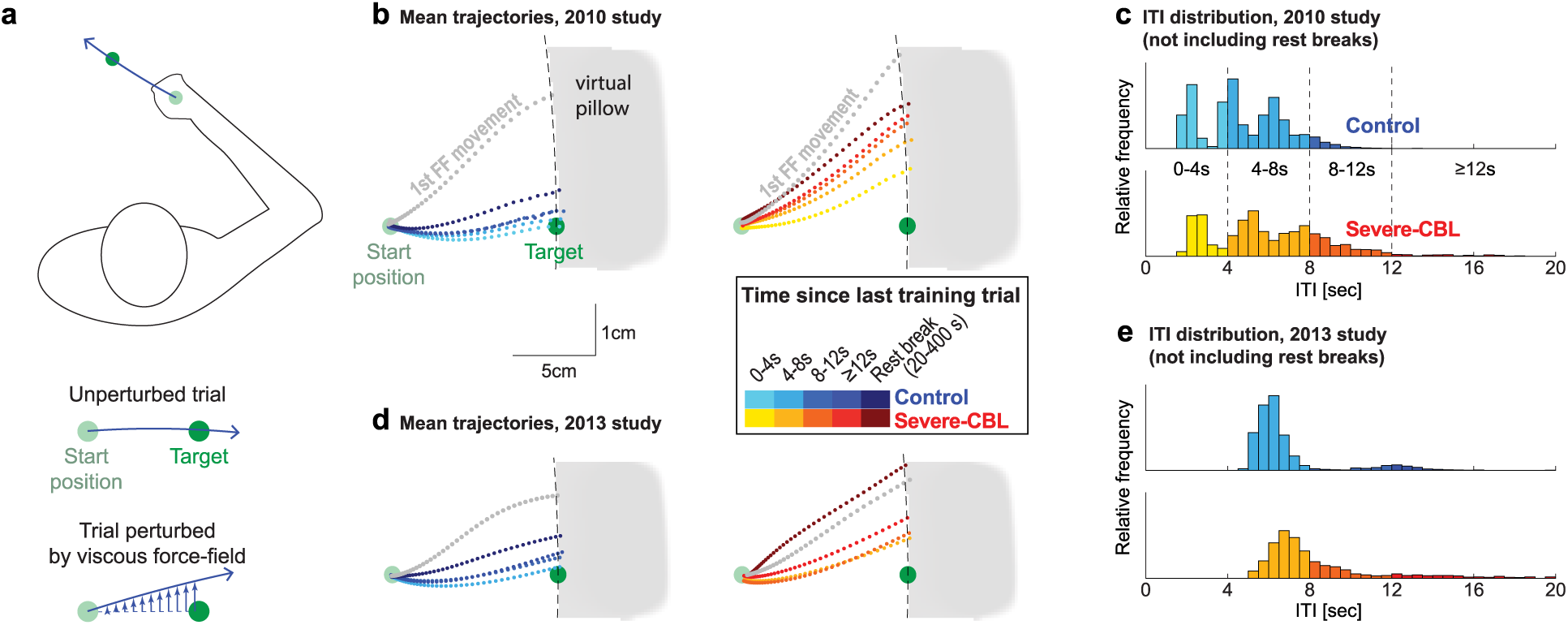
Experiment summary. **(a)** Top: top-down view of the experiment setup. Using a robotic arm, participants made reaching movements on a horizontal plane where they had to shoot-through targets displayed on a screen. Bottom: examples of movement types. During unperturbed trials, participants simply shot through the target. During training, movements were perturbed by a velocity-dependent curl force-field exerted by the robotic arm, which pushed the participant’s hand lateral to the movement direction. In both unperturbed and perturbed trials, after each movement went beyond the 10cm target, it was brought to a halt by a virtual pillow. **(b)** Average reaching trajectories in the abrupt-onset perturbation block from the 2010 study for initial exposure to the FF (gray), non-post-break trials grouped based on the preceding inter-trial interval (ITI) (blue shades for Controls, red shades for Severe-CBL), and post-break trials (dark blue for controls, dark red for Severe-CBL). Trajectories have been rotated so the movement direction is shown as rightwards in all cases. Controls are displayed on the left; participants with severe cerebellar degeneration (Severe-CBL group, ICARS≥40) on the right. Note that movement-direction errors appear more sensitive to increased ITIs for Severe-CBL participants (dark red vs yellow) than for controls (dark blue vs light blue), in line with increased decay of sensorimotor memory over the illustrated time span. This effect is examined in Figures 3-4. **(c)** Distribution of ITIs (excluding post-break trials), colored based on ITI value. Top: Controls; Bottom: Severe-CBL. Note the large spread of ITIs, which allows assessment of the sensitivity of motor memories to time in Figure 4. **(d,e)** Same as **(b,c)** but for the 2013 study.

For abrupt training, the sudden FF onset resulted in similar initial deflections of the direction of movement for both Control and Severe-CBL groups (15.2±1.3° (mean±SEM) and 17.9±3.1°, respectively, for the 2010 data; 8.3±0.7° and 8.8±1.9° for the 2013 data, the gray traces in Figure 1b,d). The approximately 2-fold difference between the initial deflections observed for the two studies was somewhat surprising given the that the size of FF perturbations was very similar (11 N/(m/s) for the 2010 study and 13 N/(m/s) for the 2013 study), but was likely due to differences in the physical inertia of the robotic arms used. We quantified the deflections elicited by the FF perturbations with the same measure used in the original studies: the movement direction error observed when the participant’s hand reached the target distance. We used the value of direction error at the onset of the abrupt FF condition to normalize subsequent reductions in it into a learning index measured for each trial in both the abrupt and gradual conditions after subtracting out the baseline movement direction (see Materials and Methods, Equation 2). This learning index would be 0 if no learning was present and 1 for full learning. Note that, for the gradual data, we further normalized the learning index by the relative FF strength so that the learning index on each trial measured performance with respect to the size of the current FF perturbation.

### Deficits in sensorimotor memory are markedly increased following longer memory windows in cerebellar degeneration

To examine sensorimotor learning that was at or near asymptote, we focused our analysis on the last three-quarters of training trials (last 180 out of 240 training trials from the 2010 study, last 129 out of 170 training trials from the 2013 study, see Figure 2). Consistent with the reports in the original manuscripts, we found that overall asymptotic learning displayed a modest yet highly significant 35±7% deficit (mean±SEM across participants) for the Severe-CBL group compared to controls (average learning index for the combined FF training data (abrupt + gradual across both studies): 0.56±0.06 for Severe-CBL vs 0.87±0.01 for controls, t(46)=5.5, p=7.8 ×10^−7^). This deficit was comparable when we separately examined the data from the 2010 study (29±4%) and the 2013 study (39±11%) and for separate analyses of the abrupt (39±6%) and gradual (31±15%) conditions.

**Figure 2:**
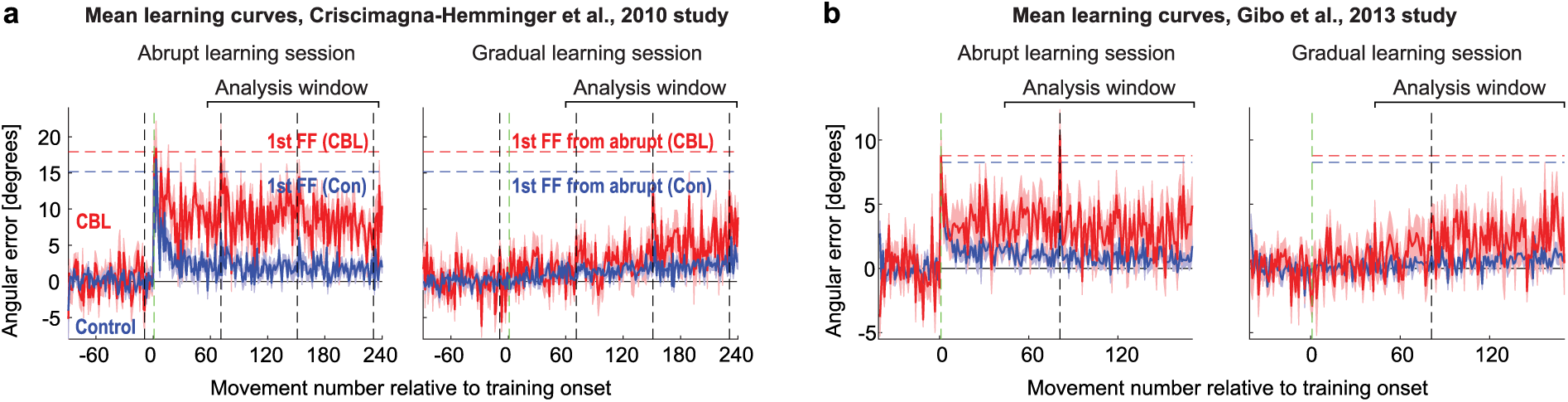
Comparison of overall learning curves. **(a)** Mean learning curves for the 2010 study, showing angular error for each trial for Controls (blue) and Severe-CBL (red). Black vertical dashed lines indicate trials following a break and the green vertical line indicates the introduction of the FF (1^st^ FF trial). Note that, because the 2010 study used two movement directions, there were two post-break trials following each rest break, and two 1^st^ FF trials; for clarity, only the first of these pairs of trials is indicated. Horizontal dashed lines indicate the average error during the 1^st^ FF trial for the abrupt data, which was used to normalize error in computing the learning index. Light blue and light red shading indicates SEM. Note the modest deficit in overall sensorimotor learning (<40%) for Severe-CBL participants, and that trials following rest breaks appear to display increased error, especially for severe-CBL patients, in line with a deficit in temporally-persistent learning, as examined in Figure 3. **(b)** Same as **(a)** but for the 2013 study.

We found, however, a considerably larger deficit when the inter-trial time interval (ITI) was high. Three rest breaks were included during both abrupt and gradual FF training periods in the 2010 study and one was included during both abrupt and gradual FF training periods in the 2013 study. The trials following these rest breaks had systematically greater ITIs than non-post-break trials (90% c.i. across all trials: 25-243s and 2-12s, respectively), so that the time windows for memory maintenance would be greater for post-break than for non-post-break trials. Specifically, in the 2010 study, average ITIs were 67.3±11.2s vs 6.7±0.5s for post-break and non-post-break trials for the Severe-CBL group (mean±SEM across participants), and 41.3±2.2s vs 5.3±0.1s for the control group. And in the 2013 study they were larger still, with average ITIs of 182.0±18.8s vs 8.4±0.4s for post-break and non-post-break trials for the Severe-CBL group, and 235.7±56.9s vs 6.8±0.1s for the control group. Critically, we found that these larger post-break ITIs were associated with a dramatic 86±19% deficit in the sensorimotor memory available during motor command generation for the Severe-CBL group vs controls (learning index values of 0.09±0.12 vs 0.65±0.08 in Severe-CBL vs Controls, t(43)=3.9, p=0.00016 overall, 0.26±0.09 vs 0.66±0.03, t(19)=4.9, p=5.2×10^−5^ in the 2010 study, and –0.05±0.20 vs 0.63±0.16, t(22)=2.7, p=0.0069 in the 2013 study; and for separate analyses of the abrupt and gradual conditions: 0.07±0.16 vs 0.63±0.06, t(39)=3.9, p=1.8×10^−4^ for the abrupt data and 0.17±0.17 vs. 0.70±0.13, t(37)=2.6, p=0.0071 for the gradual data). This 86% deficit on post-break trials is remarkable in comparison to the 35.2% deficit present in the same data set when post-break trials were not broken out or the 34.5% deficit present in non-post-break trials. The deficit on post-break trials was markedly lower for patients with milder cerebellar degeneration (Figure S1); this is in line with the observation that reduction in slow-process persistent learning can be subtle in groups with ICARS<40^58^.The disproportionally large deficit for post-break compared to non-post-break trials is in line with specific impairment in temporally-persistent, rather than temporally-volatile, sensorimotor memory. These findings indicate that even in paradigms where, overall, the majority of sensorimotor learning is preserved in individuals with severe cerebellar degeneration, sensorimotor memory can be markedly reduced when windows for memory maintenance are prolonged to half a minute or greater.

A potential limitation of the data from the 2010 study is that the post-break ITIs for the Control group were about 1/3 shorter than that for the Severe-CBL group (41.3±2.2s vs 67.3±11.2s). This was not the case in the 2013 study, as there the post-break ITIs for the Control group were instead 1/3 longer than that for the Severe-CBL group (235.7±56.9s vs 182.0±18.8s), but it still raised the possibility that the difference in post-break adaptation we observed in the data from the 2010 study might be primarily explained by the fact that Controls simply had less time to decay. To investigate this, we performed an analysis that compared Severe-CBL and Control groups when post-break ITIs were matched. Specifically, we removed the trials with the most extreme ITIs from both groups (i.e. the longest ITIs for Severe-CBL and the shortest for Controls), until the mean ITI for the Severe-CBL group crossed under that for the Control group (47.29 vs 47.35s, respectively). To achieve this, 37% of post-break trials were excluded from each group. This exclusion, however, had little effect on the mean learning index values in either group (0.29±0.10 vs 0.26±0.09 for the ITI-matched vs original data for Severe-CBL; 0.61±0.04 vs 0.66±0.03 for Controls). Correspondingly, temporally-persistent (post-break) learning index values remained significantly different between the two groups (t(19)=3.6, p=0.0011), indicating that the reduction in post-break adaptation indices observed in the data from the 2010 study were primarily due to inherent differences between the groups rather than the differences in post-break ITIs.

### Dissecting sensorimotor memory into temporally-volatile and temporally-persistent components reveals that cerebellar degeneration specifically impairs temporally-persistent memory

A temporally-volatile component of sensorimotor memory has been shown to decay exponentially with a time constant of just 20 seconds^55–57^. As this is considerably shorter than the ITIs for post-break trials, the vast majority of this memory (nearly 90% by 40sec) would have decayed away by the time that post-break trials are executed, thus allowing post-break trials to effectively isolate temporally-persistent memory. We, therefore, operationally defined the temporally-persistent component of memory as the memory available on post-break trials, and temporally-volatile memory as the difference between non-post-break and post-break trials (see Figure 3a-c).

**Figure 3:**
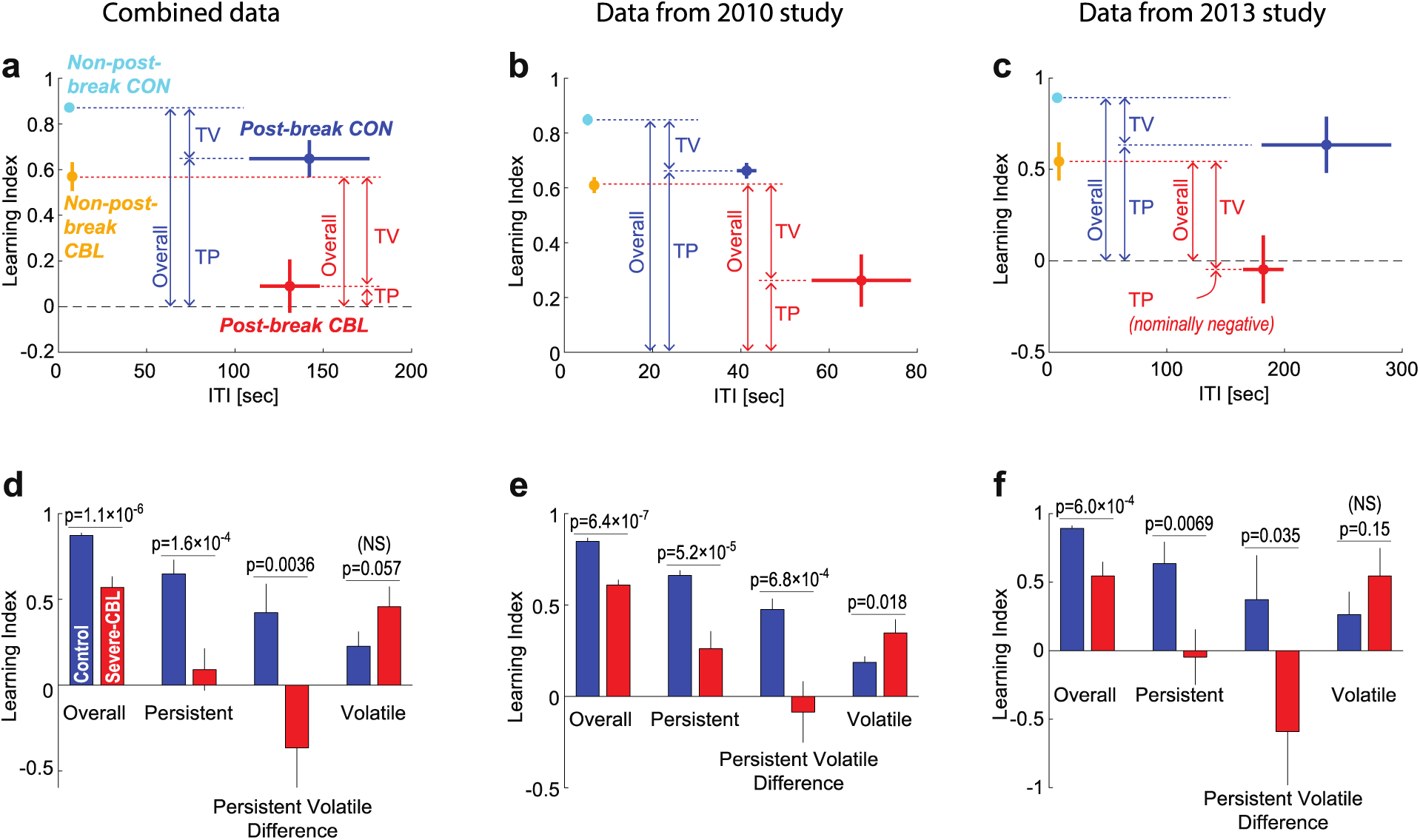
Severe-CBL participants display a specific impairment in temporally-persistent sensorimotor memory. **(a)** Mean learning index values are plotted against the corresponding average ITIs for non-post-break and post-break trials and for controls (light blue and blue, respectively) and severe-CBL participants (orange and red, respectively), for the data combined across the 2010 and 2013 studies (Severe-CBL: n=20; Controls: n=28). The short memory window observed for non-post-break trials allows both temporally-volatile (TV) and temporally-persistent (TP) sensorimotor memories to be present. In contrast, the long memory window observed for post-break trials eliminates temporally-volatile (TV) memory so that only temporally-persistent (TP) memories are present. Therefore, as illustrated by the vertical double-ended arrows, the overall memory present on non-post-break trials (TV+TP) can be dissected into individual components corresponding to the post-break memory level (TP) and the difference between the memory levels on non-post-break and post-break trials (TV). Results show only a modest impairment for overall memory, and no impairment for TV memory, but a near-complete impairment for TP memory for the severe-CBL data compared to controls. Data from both studies together; errorbars indicate SEM). **(b)** Same as **(a)** but for data from the 2010 study (Severe-CBL: n=8; Controls: n=13). **(c)** Same as **(a)** but for data from the 2013 study (Severe-CBL: n=12; Controls: n=15). **(d)** Comparison between the levels of overall memory, temporally-persistent memory, the difference between temporally-persistent and temporally-volatile memory, and temporally-volatile memory for Control vs Severe-CBL participants. Note that the difference between TP and TV memory levels, which measures the specificity of the impairment to TP memory, is not only significantly smaller in severe-CBL participants compared to controls, but also negative for Severe-CBL while positive for Controls, indicating the TP memory dominates TV for Controls while TV memory dominates TP for Severe-CBL participants. **(e)** and **(f)**: Same as **(d)** but for the 2010 and 2013 studies, respectively.

Our hypothesis that cerebellar degeneration specifically impairs longer-term, temporally-persistent memories rather than short-term, temporally-volatile memories, predicts that the balance between temporally-persistent and temporally-volatile memory will shift towards the latter for Severe-CBL individuals relative to Controls. This shift would reduce the difference between the levels of temporally-persistent and temporally-volatile sensorimotor memory for the Severe-CBL group compared to Controls. We, therefore, tested this prediction by calculating the difference between temporally-persistent and temporally-volatile sensorimotor memory for each individual and then comparing the results for the Severe-CBL vs Control groups. In agreement with the prediction, we found a significant reduction of this difference for Severe-CBL participants compared to Controls (−0.37±0.23 vs +0.42±0.17, t(43)=2.8, p=0.0036 overall, Figure 3d; –0.09±0.17 vs. +0.48±0.06, t(19)=3.8, p=0.00068 for the 2010 study [Figure 3e] and –0.59±0.39 vs. +0.37±0.32, t(22)=1.9, p=0.035 for the 2013 study [Figure 3f] separately; –0.38±0.30 vs. +0.39±0.11, t(39)=2.8, p=0.0036 for the abrupt and –0.29±0.25 vs. +0.55±0.25, t(37)=2.3, p=0.0142 for the gradual data separately), indicating a significantly greater deficit in temporally-persistent than temporally-volatile learning. The *positive* values consistently observed for Controls indicate *greater* levels of persistent than volatile memories, in contrast to the *negative* values consistently observed for Severe-CBL, which indicate that these individuals instead displayed *smaller* levels of persistent than volatile memories. This finding demonstrates a dissociation between the effects of cerebellar degeneration on the ability to form temporally-persistent and temporally-volatile memories, indicating a specific deficit in temporally-persistent learning.

The prediction that the balance between temporally-persistent and temporally-volatile memories will shift towards the latter in the Severe-CBL group can be independently tested by an analysis of trial-to-trial differences within the non-post-break data, where ITIs were small compared to the post-break data. If Severe-CBL individuals indeed display a higher proportion of temporally-volatile learning (which is prone to decay with time) vs temporally-persistent learning (which is not), then the overall memory would display greater decay with time. And although the ITIs in these data, being smaller than the time constant reported for the decay of temporally-volatile memory, should result in only partial temporally-volatile decay, the amount of this decay per unit time can be measured based on the ITI variability existent in the within-block data (Figure 1c,e). In the 2010 study, this ITI variability primarily arises from the random presentation of one of the two possible targets on each trial. Thus, the same target could be visited on consecutive trials resulting in a low same-target ITI, or could be visited after multiple intervening movements to the other target, resulting in a high same-target ITI. Note that it is the same-target rather than the target-independent ITI that should be relevant for memory retention here, as the two targets were 180° apart and sensorimotor memories formed for movements 180° apart are distinct, with limited transfer of learning from one direction to the other^57,59–61^. In the 2013 study, there was only a single target location, reducing the spread of ITIs, with the presence of occasional (10% of total learning trials) single error-clamp trials (in which the error was clamped to zero, thus preventing error-based learning) adding only a small amount of variability to the random ITI fluctuations that occurred during the experiment. Consequently, the average variance of the within-block ITI was greater in the 2010 study compared to the 2013 one (for Severe-CBL: 12.0 sec^2^ vs. 6.8 sec^2^; for Controls: 7.0 sec^2^ vs. 4.0 sec^2^), but the effects of this variability on the availability of sensorimotor memory can be nevertheless examined in both cases.

To test our prediction of steeper decay in patients with cerebellar degeneration, we estimated the rate of time-dependent decay within the non-post-break data by regressing the relative learning index on each trial onto the corresponding ITI for each participant (see Materials and Methods for details). We found regression slopes to be significantly different in the Severe-CBL vs. Control groups (Figure 4a, decay rates of 0.036±0.008 vs. 0.018±0.003 learning index units (which are dimensionless) per second, t(46)=2.6, p=0.0066 for the combined data, with 0.027±0.005 vs. 0.015±0.003 sec^−1^, t(19)=2.4, p=0.014 for the 2010 study [Figure 4b] and 0.042±0.012 vs. 0.020±0.004 sec^−1^, t(25)=1.8, p=0.039 for the 2013 study [Figure 4c]). This finding is in line with a shift towards a temporally-volatile memory of learning in patients with cerebellar degeneration, consistent with an impairment in temporally-persistent memory.

**Figure 4:**
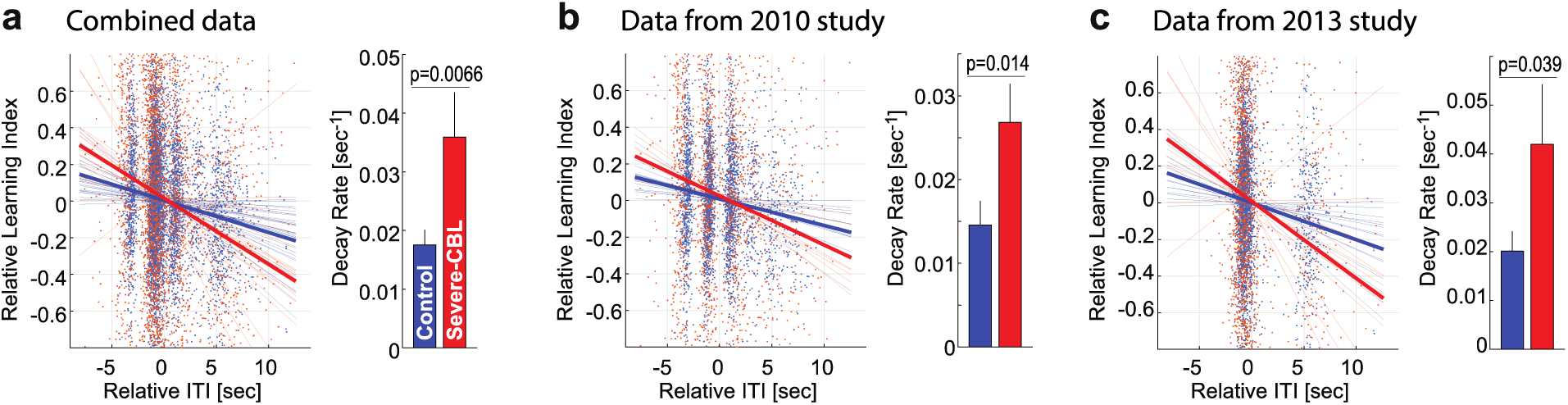
Severe-CBL participants display greater temporal decay of sensorimotor memory. **(a)** Left: Relationship between learning index and ITI within non-post-break trials. The y-axis shows the relative learning index for each trial (normalized and mean subtracted), whereas the x-axis shows the corresponding relative ITI. Thick lines indicate average fits for each group, whereas thin lines indicate individual fits. Right: Bars show the sensitivity (slope) for the relationship plotted at left. Severe-CBL participants display decay rates that are 2x faster than Controls. **(b)** and **(c)**: Same as **(a)** but separately for the 2010 and 2013 studies, respectively.

Our hypothesis of a specific impairment of longer-term temporally-persistent memory in cerebellar degeneration makes a somewhat counterintuitive prediction for short-term temporally-volatile memory. It predicts that individuals with cerebellar degeneration would show not only preserved levels of short-term temporally-volatile memory that contrasts with reductions in longer-term temporally-persistent memory, but beyond that, it predicts levels of temporally-volatile memory that would be even greater than those found in healthy individuals during sensorimotor learning. This counterintuitive increase in temporally-volatile memory would occur because impaired temporally-persistent memory should result in larger motor errors which would provide greater drive for temporally-volatile learning, leading to increased levels of temporally-volatile memory. If the specificity of the impairment in temporally-persistent memory was sufficiently profound that the effect of this increased drive consistently outweighed any non-specific impairment in short-term temporally volatile learning, then systematically higher levels of temporally-volatile memory should be present in Severe-CBL patients compared to controls. We tested this prediction by comparing the levels of temporally-volatile sensorimotor learning between the Severe-CBL group and controls. Remarkably, the Severe-CBL group displayed a level of temporally-volatile memory that was not lower but rather nominally 2-fold higher than the control group, in line with the prediction (Figure 3d-f, learning index values of 0.46±0.12 vs. 0.23±0.09 for Severe-CBL vs. Controls, t(43)=1.6, p=0.057; 0.35±0.07 vs. 0.19±0.03, t(19)=2.3, p=0.018 for the 2010 study and 0.54±0.21 vs. 0.26±0.17, t(22)=1.1, p=0.15 for the 2013 study separately). The nominal increase in temporally-volatile learning observed in the combined data when comparing Severe-CBL vs Control participants is in line with the hypothesized increase in the level of temporally-volatile sensorimotor memory. But with significant effects only observed in the data from the 2010 study, the current data do not provide reliable evidence for it.

### Analysis of the findings from other sensorimotor learning studies

The specific deficit in temporally-persistent sensorimotor learning that we observed in the detailed trial-by-trial analysis of the data from the Gibo et al., 2013 and Criscimagna-Hemminger et al., 2010 studies manifested as increased impairment in individuals with cerebellar damage as ITIs were increased. This would predict that studies that employ systematically longer ITIs would report systematically greater levels of impairment than those with shorter ITIs. Unfortunately, information about ITIs is rarely reported in studies of sensorimotor learning. However, studies often differ in the number of movement directions present in the training paradigm, which is consistently reported. And paradigms with a greater number of movement directions should display consistently longer intervals between consecutive trials in the same direction if movement directions are either randomly sequenced or cycled, as is the case for all relevant studies we identified. Because movement directions separated by more than 30° or 45° are learned largely independently^57,59,60^, it is this within-direction ITI that is relevant for sensorimotor memory. Thus, our hypothesis would predict that studies with a greater number of well-separated movement directions (separated by 45° or more) would display consistently higher levels of impairment than those with a smaller number. The evidence supporting this prediction would be muddied, of course, by any differences in the severity of ataxia in the populations studied, and the inexact relationship between the number of movement directions and within-direction ITIs. However, the effects of these issues would be mitigated if multiple studies with large and small numbers of movement directions could be identified.

Beyond the two analyzed above^49,50^, we were able to identify 15 published studies that compared sensorimotor learning for upper limb reaching movements between patients with cerebellar ataxia and healthy controls^41–46,48,51,52,62–67^. Comparisons of results across these studies reveal a clear pattern whereby a larger number of movement directions was consistently associated with greater impairment. Studies using 4-8 movement directions, in both FF^51,52^ and visuomotor rotation tasks^44–46,62–64^, reported deficits of 56% on average in the amount of sensorimotor learning displayed by patients with cerebellar damage relative to controls (range 28-76%), with only one study displaying a deficit below 45%^62^. In comparison, studies using 1-3 movement directions reported deficits that were nearly 3-fold lower (22% on average, range 2-43%; t(13)=4.4, p=0.00075 for comparison of deficits between these two groups of studies)^41–43,48,65–67^. Overall, we find that the average difference between the deficits observed in these two conditions explains a remarkable 60% of the variance across these 15 previous studies. That this is the case even though (a) the link between the number of movement directions and mean within-direction ITIs is indirect, and (b) differences in severity of cerebellar ataxia are not adjusted for, underscores the strength of the effect.

## Discussion

Here we hypothesized that cerebellar damage impairs the formation of temporally-persistent memory for sensorimotor learning, leaving short-term, temporally-volatile memories intact. To test this hypothesis, we reexamined data from two previous studies that had reported only modest deficits in sensorimotor learning for individuals who displayed severe ataxia from cerebellar degeneration, with the prediction that more profound deficits would be present on individual trials that required memory maintenance beyond the time window for temporally-volatile memory.

Our primary analysis, therefore, was a dissection of memory from sensorimotor learning into temporally-persistent and temporally-volatile components based on the prolonged inter-trial time intervals (ITIs) that occurred during the occasional rest breaks that were built into these experiments. We found the amount of sensorimotor memory remaining after these prolonged ITIs to be dramatically decreased with severe cerebellar damage: Temporally-persistent learning measured on post-break trials was *almost absent* in patients, with a remarkable 86% deficit compared to controls, in stark contrast to the modest 34% deficit observed on non-post-break trials where temporally-volatile learning could also contribute to sensorimotor memory. This finding indicates dramatically impaired temporally-persistent sensorimotor memory in individuals with severe cerebellar ataxia and suggests an impairment specific to temporally-persistent memory as the deficit is mitigated on non-post-break trials.

When we directly tested the specificity of the impairment across temporally-persistent and temporally-volatile sensorimotor memories by comparing the difference between them, we found significantly greater impairment for temporally-persistent than temporally-volatile memory in the severe CBL group (p<0.002). A significant difference was observed not only in the combined data, but also independently in each dataset (2010 vs 2013 study) and in each learning condition (abrupt vs gradual training). In fact, the difference we observed was so large that the memory measured on non-post-break trials where temporally-persistent and temporally-volatile memories could both be present, was dominated by temporally-persistent memory in healthy controls but flipped to be dominated by temporally-volatile memory in the severe-CBL group in the combined data as well as in each individual study (see Figure 3d-f). These findings provide clear evidence for the prediction of a specific deficit in temporally-persistent sensorimotor memory in severe CBL participants.

Two secondary analyses further examine the degree of this specificity. First, we reasoned that the shift in the composition of sensorimotor memory between temporally-persistent and temporally-volatile constituents resulting from a specific deficit in temporally-persistent memory should lead to a more rapid memory decay in the non-post-break data. We therefore regressed sensorimotor learning in non-post-break trials against the duration of the memory window to estimate the rate of memory decay. This analysis revealed memory decay that is more rapid in severe CBL participants in line with a shift toward greater temporally-volatile learning. Second, we found that the level of temporally-volatile memory itself was not only unimpaired but paradoxically increased in severe CBL participants, an effect that would be predicted by a deficit in temporally-persistent memory that is so specific that the increase in motor error arising this deficit would increase the drive for temporally-volatile learning by an amount that outweighs any reduction in temporally-volatile learning ability.

Our findings were further supported by analysis of the deficits reported in 15 other studies of sensorimotor adaptation in patients with cerebellar damage. While detailed inter-trial timing information was not available to us, studies which used a larger number of target directions (4-8), which would result in a long average ITI between movements to the same target, reported 3-fold greater deficits, on average, compared to studies using fewer movement directions (1-3).

Taken together, these findings advance our understanding of the role of the cerebellum in sensorimotor learning, identifying a highly specific contribution to temporally-persistent sensorimotor memory that is dramatically impaired in individuals with severe cerebellar ataxia. Given previous work showing that temporally-persistent learning leads to long-term retention^68,69^, our results point toward a role for the cerebellum as a gateway to the formation of stable procedural memories, akin to the well-studied role of the MTL as a gateway to the formation of stable declarative memories. And they suggest that, like declarative learning, sensorimotor learning engages largely independent short-term and long-term memory systems.

Note that here we assessed the decay of sensorimotor memory as a reduction in the learning index – that is, the degree to which mean motor behavior reverted back to baseline. Measuring memory decay based on mean effects is typical in the sensorimotor learning literature^56,57,70^; however, memory decay can be also assessed based on decreased precision of the memory^71^. Since differences in precision admit differences in certainty, these seemingly different approaches would be tied together in a Bayesian framework that would predict that a decrease in the certainty about a sensorimotor memory would result in a posterior estimate that decays closer to baseline^72^.

### Role of explicit adaptation

It is well-known that sensorimotor learning can be comprised of implicit and strategic contributions^73–76^. It is, thus, conceivable that the dichotomy between implicit and strategic contributions could at least partially explain the dichotomy we uncover between reduced temporally-persistent and spared or increased temporally-volatile memory in severe cerebellar degeneration. In particular, if strategic contributions were specific to temporally-volatile memory and were unimpaired in severe cerebellar degeneration, but implicit contributions were uniformly impaired across both temporally-volatile and temporally-persistent memories, the predicted pattern of results would be similar to what we observed. However, this scenario is implausible for multiple reasons. First, the perturbations employed in the studies we analyzed (measured by the errors observed in the first abrupt force-field exposure) were rather small (Figure 2): 15-18° in the 2010 study and 8-9° in the 2013 study. But previous studies have shown that small 15° perturbations result in minimal strategic contributions^75,77^, suggesting that high levels of strategy use were unlikely. Even when perturbations are larger, which allows explicit strategies to be readily elicited during visuomotor adaptation^77–79^, participants fail to be able to consistently invoke strategies to improve performance in FF adaptation tasks when different FFs are explicitly cued^80–83^. And even when participants are explicitly instructed to use a strategy when only a single FF is present, strategy use makes only a small contribution to the overall adaptation^76^. Second, even if high levels of strategy use were somehow present in the data we examined, strategic contributions to sensorimotor learning show a significant impairment of about 40% in cerebellar degeneration when individuals must devise their own strategies^44^, as would have to be the case for the studies we analyzed. But this is at odds with the unimpaired (and nominally 2-fold increased) temporally-volatile memory we observe (Figure 3d-f), suggesting that this unimpaired TV memory cannot be strategic. Third, even if it were somehow the case that strategy was present without the 40% deficit that has been observed in CBL patients, the expected effects would still be inconsistent with our results. This is because strategy use invariably results in extremely high adaptation levels, often 95-100%, even if implicit learning is low^73,75,84^, but this is clearly at odds with our severe-CBL data, where the learning index is 56%. Fourth, strategic contributions have been found to be temporally-persistent rather than temporally-volatile^56,85,86^, but this would prevent strategy learning from explaining the unimpaired temporally-volatile learning and thus the volatile vs persistent dichotomy that we observe. Fifth and finally, gradual training strongly reduces strategy use^44,87^; however, the dichotomy we observe between temporally-volatile and temporally-persistent learning is essentially identical between abrupt training blocks and gradual blocks, suggesting that little strategy use was present. Together, these issues make it unlikely that strategy use makes a significant contribution to the dichotomy we observe between temporally-volatile and temporally-persistent learning.

### Interaction between the timescales of sensorimotor memory and computational function in the cerebellum

Our findings show that the cerebellum is required for the formation of temporally-persistent sensorimotor memories which lead to long-term memory formation^68,69^ but not for short-term temporally-volatile memories. While the timescales of cerebellar contributions to sensorimotor memories are a distinct issue from the nature of the neural computations performed by cerebellar circuits that support sensorimotor control, our findings do suggest that these computations have a specific influence on long-term vs. short-term sensorimotor memories. Although not fully understood, the neural computations in cerebellar circuits are widely believed to support internal predictions^35,36,88,89^, and thus our findings suggest that predictive information is more likely to be present in longer-term sensorimotor memories than short-term, temporally-volatile memories.

### Parallels between the cerebellum and the medial temporal lobe

Our results provide compelling evidence for the idea that, like the MTL, cerebellar damage has a greater effect on the formation of long-term than short-term memory, albeit for sensorimotor memories rather than declarative ones. In fact, the current evidence (see Figure 3) suggests that the dichotomy between the effects of cerebellar damage on long-term vs short-term memory is essentially complete, and perhaps even more extreme than the corresponding dichotomy for the effect of MTL damage on long-term vs short-term memory, which is likely incomplete based on recent work^6,7,9,12^.

The functional parallel we find here between the MTL and the cerebellum – that these two structures may act as parallel gateways to the formation of long-term declarative and sensorimotor memories – is further supported by highly regular neural architecture in both structures and their connectivity patterns. In line with its function as a gateway to the formation of long-term declarative memories, the MTL has widespread reciprocal connections with the cortex where these memories eventually reside^13,90–93^. Correspondingly, neuroanatomical and functional studies have both shown widespread reciprocal connections between the cerebellum and cortical areas involved in motor planning and execution, including M1, premotor cortex, and parietal cortex, but also to cortical areas related to executive function and decision making^94–98^.

Whereas MTL function is primarily known for playing a greater role for long-term compared to short-term memory, it is also known to play a greater role in the acquisition compared to the storage of long-term memory, as the MTL is required for the acquisition of long-term declarative memory but not its eventual storage^13,99^. Intriguingly, previous work provides evidence that the cerebellum might analogously be required for the acquisition of sensorimotor memory but not its eventual storage^100^. In particular, cerebellar damage in humans dramatically impairs the ability to acquire new sensorimotor memories but preserves previously-learned sensorimotor memories both for anticipatory postural adjustments (APAs)^101^ and eyeblink conditioning^102–104^.

We note, however, that there is evidence in non-human mammals that different regions of the cerebellum might play different roles in acquisition and long-term storage of sensorimotor memory, with the cerebellar cortex being required for the acquisition but not the eventual storage of optokinetic responses^105^, analogous to the role of the MTL for declarative memory, but deep nuclei in the cerebellum being required for both acquisition and storage of optokinetic responses and eyeblink conditioning^105–107^. Whether the apparent differences, in terms of the role of deep cerebellar nuclei, between the human and animal studies are due to inter-species differences, task differences, the ability to exert fine-grained spatial control over lesions in animal models, or another factor is not understood. However, in sum, the evidence suggests that structures in the cerebellum, either the entire cerebellum or the cerebellar cortex, can play a transient role in learning specific to the acquisition of sensorimotor memory, with long-term storage either in deep cerebellar nuclei or outside of the cerebellum altogether.

The current findings do not suggest that the cerebellum completely complements the MTL by providing a gateway for *all* long-term non-declarative memories. In fact, evidence suggests that several types of non-motor procedural learning do not strongly depend on the cerebellum. Patients with cerebellar damage appear to be unimpaired in tasks such as implicitly learning artificial grammars^108^, making probabilistic associations^109^, or implicit category learning^110^. On the other hand, cerebellar function is not limited to supporting sensorimotor learning, as the cerebellum has widespread connections to areas not directly involved in sensorimotor function^111–115^, and cerebellar damage impairs executive function, spatial cognition, and language^116–118^. Further work is needed to delineate the types of procedural learning that are dependent on the gateway function of the cerebellum we identify here.

## Methods

### Participants

No new participants were recruited for the present study; instead, we reanalyzed data from ^49^ and ^50^. Data from a total of n=20 patients with severe cerebellar ataxia (Severe-CBL group, ICARS score ≥40 – 2010 study: n=8, ICARS=53.1±5.7 [mean±std]; 4 male, 57.5±14.0 y.o.; 2013 study: n=12, ICARS=54.5±11.8; 8 male, 57.6±10.4 y.o.) and n=28 healthy controls (Control group – 2010 study: n=13, 58.5±11.4 y.o.; 2013 study: n=15, 58.7±9.7 y.o.) are included in the main analysis. These patients were diagnosed with either a genetically identified spinocerebellar ataxia subtype (SCA6, n=7, SCA8, n=2, SCA14, n=1, and a single individual with both SCA6 and SCA8), autosomal dominant cerebellar ataxia that was not genetically identified (ADCA; n=4), or a sporadic case of ataxia (n=5). Four of the patients with severe cerebellar ataxia and six of the controls were left-handed; participants performed the task with their dominant hand (with the exception of one left-handed patient). An additional 8 patients with milder cerebellar ataxia (Milder-CBL group, ICARS score <40 – 2010 study: N=5, ICARS=24.8±11.9, all female, 60.0±6.6y.o.; 2013 study: N=3, ICARS=21.0±9.5, 1 male, 63.0±5.3 y.o., 1 left-handed) are examined in the supplementary analysis.

Importantly, both studies included only individuals with purely cerebellar signs based on examination of upper limb function which was used on these tasks. In particular, the presence of extra-cerebellar signs including hypertonia, proprioceptive loss or reduced sensitivity to fine touch were used as exclusionary criteria^49,50^. SCA6 is believed to be a purely cerebellar syndrome, based on evidence from neuroimaging and neuropathological examination finding atrophy confined to the cerebellum^119,120^; similarly, imaging in SCA8 suggests atrophy limited to the cerebellum, with only minimal brainstem involvement^121^. For patients with SCA14, imaging and neuropathology studies also show involvement that is generally exclusive to the cerebellum^122–127^.

### Tasks

Detailed descriptions of the tasks for the (Criscimagna-Hemminger, et al, 2010 and Gibo et al 2013 studies that we analyzed are described therein. For convenience, we provide brief summaries below.

Both tasks involved serially-repeated goal-directed reaching arm movements that were, in the training block, physically perturbed by a velocity-dependent curl force field (FF) environment that was either gradually or abruptly introduced. This FF produced a perturbing force (*F*) lateral to the direction of motion based on the motion’s velocity (*v*):

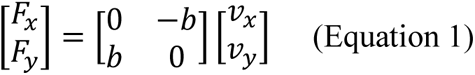

The strength of the viscous FF, *b*, employed in these studies was similar: 11 Ns/m in Criscimagna-Hemminger et al (2010), and 13 Ns/m in Gibo et al (2013). For the gradually introduced FF, the value of *b* linearly increased to its full strength over the course of training. Specifically, in the 2010 study, it gradually increased for 232 trials (116 per movement direction) and then maintained at full strength for a further 8 trials. In the 2013 study, it gradually increased for 160 trials and then maintained at full strength for a further 10 trials.

A key difference between the two studies’ design was that the 2010 study used two target locations (at 31.5° and 211.5°, i.e. close to 2 and 8 o’clock, respectively) whereas the 2013 study used one (at 0°, i.e. 3 o’clock; note that, in both studies, target locations were mirrored across the vertical axis for participants using their left arm).

## Data analysis

### Learning index

To assess adaptation to the FF for both the Criscimagna-Hemminger et al., 2010 and Gibo et al., 2013 studies, we first calculated angular error, θ_*n*_, at each trial *n*. We defined this error as the difference between the ideal movement direction (directly through the center of the target) relative to hand position at movement onset and the actual movement direction, calculated the moment the hand crossed the target distance. Movement onset was defined as the moment the tangential velocity of the hand first exceeded 5cm/sec. Specifically, beginning from the movement mid-point (when the hand was half the target distance away from the starting position), we went backwards in time until the moment velocity first dropped below 5cm/s and remained so for at least 100ms.

Because the center-out movements were relatively rapid (average movement times of 349±11ms for Severe-CBL patients and 322±11ms for Controls across both studies) and importantly, did not require stopping at the target which would make feedback corrections necessary, the errors we observe predominantly reflect the inability of a feedforward controller to counter the imposed perturbation.

Importantly, because patients may have different directional biases at baseline^50^ that could add up to their FF response, we subtracted baseline biases separately for each target and session. Following baseline subtraction, we flipped (i) CCW data and (ii) left-hand data in order to combine sessions using either hand and either FF polarity.

We then defined the learning index at each trial, *a*_*n*_, as the fractional reduction in angular error relative to the angular error at first exposure to the perturbation for the abrupt data, θ*_init_*:

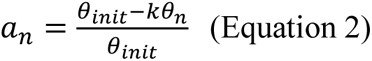

Because there were very few measurements of the denominator, θ*_init_* (only one per movement direction per participant, that is two measurements in the Criscimagna-Hemminger et al. dataset and one the Gibo et al. dataset for each participant, leading to noisy per-participant estimates) we used the average value of θ*_init_* across all participants in each group (Control, Severe-CBL, or Milder-CBL) in each dataset (Criscimagna-Hemminger et al. or Gibo et al.) prior to calculating *a*_*n*_. Importantly, we did not calculate θ*_init_* for the gradual data (as the strength of the FF for the first training trial would be very small [<1% compared to that in the abrupt case] and thus the estimation of any resulting angular errors would be overwhelmed by noise), and instead used the same values as for the abrupt data. To estimate *a*_*n*_ for gradual data, we normalized the corresponding gradual errors θ*_n_* by the relative FF strength *k* (for example, if FF strength at trial *n* were half the full strength on which θ*_init_* was based, the error would be multiplied by two). For the abrupt data, the value of *k* was 1 throughout.

In the gradual data, estimated errors for low-FF trials would suffer from a lower signal to noise ratio compared to errors for high-FF trials. This is because the strength of the signal (responses driven by the FF) would scale with the FF but the motor noise would be the same. To account for this, when averaging learning indices for the gradual data across trials (see *Interblock analysis* below), we weighted each trial by the relative force-field strength (*k*) associated with it. Moreover, when estimating regression slopes (see *Intrablock analysis* below) for normalized gradual data, we used regression which weighted square errors for each trial by the relative FF magnitude (*k*) of that trial.

A fraction of trials during the training period were error-clamp trials (20% of trials in the 2010 study and 10% of trials in the 2013 study). During error-clamp trials, the FF was turned off and, instead, the robotic arm constrained movement to a straight-line path to the target. Consequently, lateral displacements on those trials do not measure learning; and our analysis was thus based solely on FF trials. While lateral forces observed on those trials could be used to measure learning, there were no post-break error-clamp trials (except in 4 out of the 62 sessions in the 2013 study), preventing such a measurement from being dissected into temporally-persistent and temporally-volatile components.

Reported variables (overall adaptation, temporally-persistent adaptation, temporally-volatile adaptation, sensitivity of learning index to trial-to-trial differences in ITI) were calculated separately for abrupt and gradual data and then averaged to yield participant-specific estimates. One participant in the Gibo et al., 2013 dataset performed the study twice (resulting in two sessions of abrupt and two sessions of gradual training, with the polarity of the FF balanced). Since our statistical testing was based on comparisons across individuals, we averaged the results from these two sessions prior to statistical testing. This participant was from the Severe-CBL group.

### Data inclusion criteria

We excluded some movements from the analysis window based on kinematic characteristics that would lead to uncertainty about using movement direction to measure sensorimotor learning (12.4±3.6% from the Severe-CBL data and 12.7±3.4% from the Control data from the CH et al study, and 18.3±4.8% from the Severe-CBL data and 5.7±1.7% from the Control data from the Gibo et al study). Specifically, we excluded: (a) trials for which radial velocity dropped below 25% of its maximum before the target distance was reached (3.5% for Severe-CBL and 10.3% for Controls in the 2010 data; 2.2% and 1.7%, correspondingly in the 2013 data), as this was at odds with task instructions to shoot through the target; (b) trials that took >500ms after movement onset to reach the target distance or for which the hand had traveled < 1/3 of target distance by 250ms, as motor errors might be muted by feedback corrections (an additional 8.4% for Severe-CBL and 2.2% for Controls in the 2010 data; 15.2% and 4.0% correspondingly in the 2013 data); (c) trials for which the hand was more than 1.5cm away from the starting position when movement onset was detected, as these could not be readily compared to most other trials (an additional 0.17% for Severe-CBL and 0% for Controls in the 2010 data; 0.90% and 0.09% correspondingly in the 2013 data); (d) trials for which the directional error 250ms after movement onset was >90°, indicating that the movement started in entirely the wrong direction (an additional 0.30% for Severe-CBL and 0.19% for Controls in the 2010 data; none in the 2013 data). These cases, which were isolated to the 2010 data where two targets were used, consisted of movements that were mistakenly directed to the unselected target (in fact, movement directions for these trials were all within 20° of the unselected target’s direction). Additionally, experimenter notes from the Gibo et al., 2013 study indicated that some trials were excluded from the analysis in the original paper (0.92% for Severe-CBL and 0.05% for Control data within the analysis window). And, in the Criscimagna-Hemminger et al., 2010 study, movement recording started too late to capture movement onset for a small number of trials (0.22% in Severe-CBL and 0.16% in Controls). Thus, these trials were also excluded prior to further analysis.

### Definition of inter-trial time interval (ITI)

We defined the ITI preceding a given trial as the time between the onset of that trial and the onset of the first preceding trial that was both on the same direction and was not an error-clamp trial.

### Interblock analysis

For each participant, we defined temporally-persistent adaptation as the average adaptation for trials following rest breaks (in the case of the 2010 study, which used two target directions, that meant taking the post-break trials for each target direction separately). The 2010 study included three breaks during the training period (with the exception of one individual session from the Severe-CBL data and one individual session from the Control data, for which timing data indicated one additional break), whereas the 2013 study included one break during the training period. In the rare cases (only in 4 individual sessions of the 2013 study) where the trial immediately following a break was an error-clamp trial, the following non-error-clamp trial was used. We defined overall adaptation as average learning index for all FF learning trials within the analysis window that did not follow rest breaks. Finally, we defined temporally-volatile adaptation as the difference between overall and temporally-persistent adaptation.

### Intrablock analysis

We assessed the rate at which sensorimotor learning decayed by regressing learning index measured for each movement against the ITI for that movement for all intra-block FF learning trials (all trials within the analysis window that did not follow a rest break) using linear regression. To examine this rate of decay relative to overall learning for each group, we first normalized the learning indices by the average intra-block learning index separately for severe cerebellar patients and controls. We then estimated the sensitivity (slope) of this normalized adaptation indices relative to ITI for each individual.

### Statistical comparisons

To compare adaptation indices between patients and controls we used single-tailed t-tests, with the prior hypothesis that patients’ adaptation indices are lower due to their impairment. We also used single-tailed t-tests to examine differences in the temporally-persistent vs. temporally-volatile components of adaptation, with the prior hypothesis that impairment specifically targets temporally-persistent adaptation, meaning that temporally-persistent adaptation in patients would be lower than controls but temporally-volatile adaptation would be higher.

To assess impairment – the ratio of the learning index of patients over controls – we used a bootstrapping procedure which allowed us to estimate both the average impairment but also the SEM associated with it.

## Acknowledgements

We would like to thank Sarah Hemminger, David Herzfeld, Reza Shadmehr, and Amy Bastian for helping us acquire the data and Tanvi Ranjan for helpful suggestions. Support for this work was provided by the McKnight Scholar Award to MAS, a Sloan Research Fellowship to MAS, and a grant from NIA (R01 AG041878) to MAS.

## Author contributions

Design of analyses: A.M.H., M.A.S.; implementation of analyses: A.M.H.; manuscript preparation: A.M.H., M.A.S., T.L.G.; Funding: M.A.S.

## Data availability statement

Pending approval from the institutions where the data were originally collected, we will make de-identified data available on a public repository ahead of publication.

## Supplementary materials

**Figure S1:**
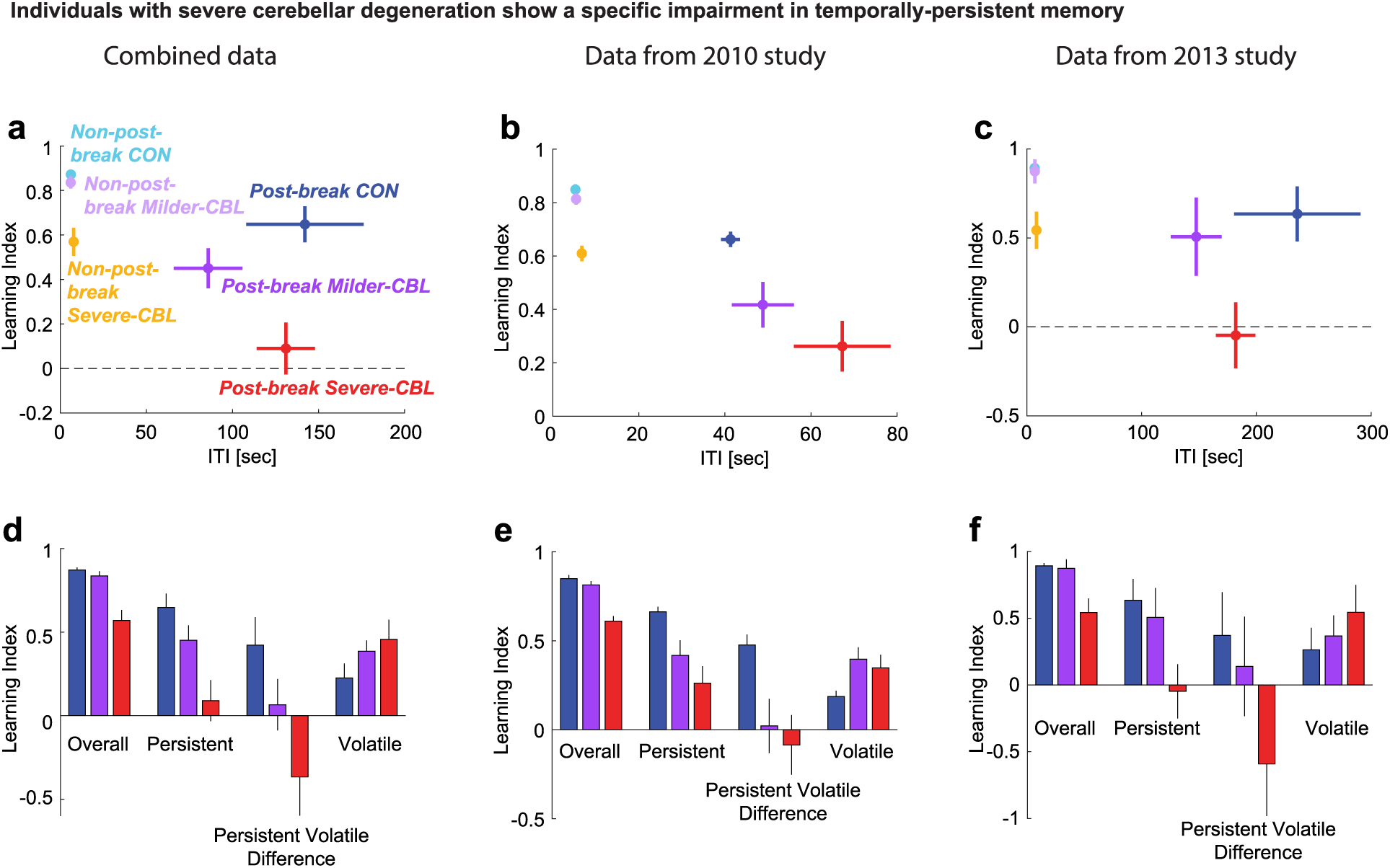
Analysis of specific impairments in temporally-persistent and temporally-volatile sensorimotor memory – including patients with milder cerebellar degeneration. Format as in Figure 3 from the main paper, but also including patients with milder cerebellar degeneration (Milder-CBL group, ICARS<40; n=5 in the 2010 study and n=3 in the 2013 study; shown in purple).

**Figure S2:**
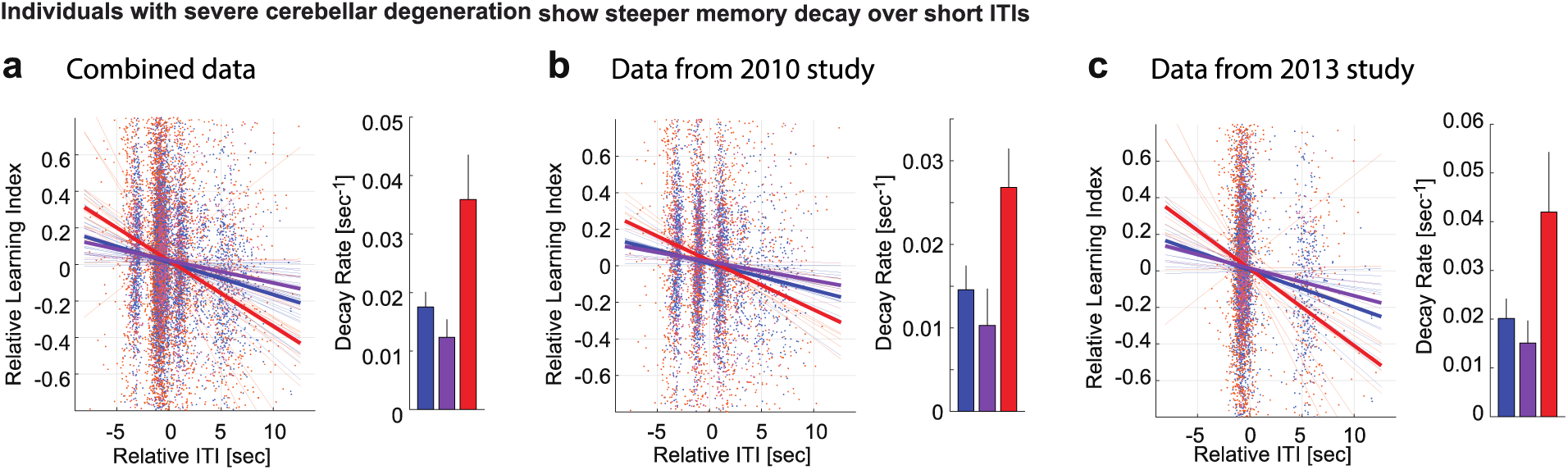
Analysis of temporal decay rates for sensorimotor memory – including patients with milder cerebellar degeneration. Format as in Figure 4 from the main paper, but also including patients with milder cerebellar degeneration (Milder-CBL group, ICARS<40; n=5 in the 2010 study and n=3 in the 2013 study; shown in purple).

