## Supplementary Figures 1 and 2 for "The cerebellum acts as the analog to the medial temporal lobe for sensorimotor memory"

### Supplementary materials

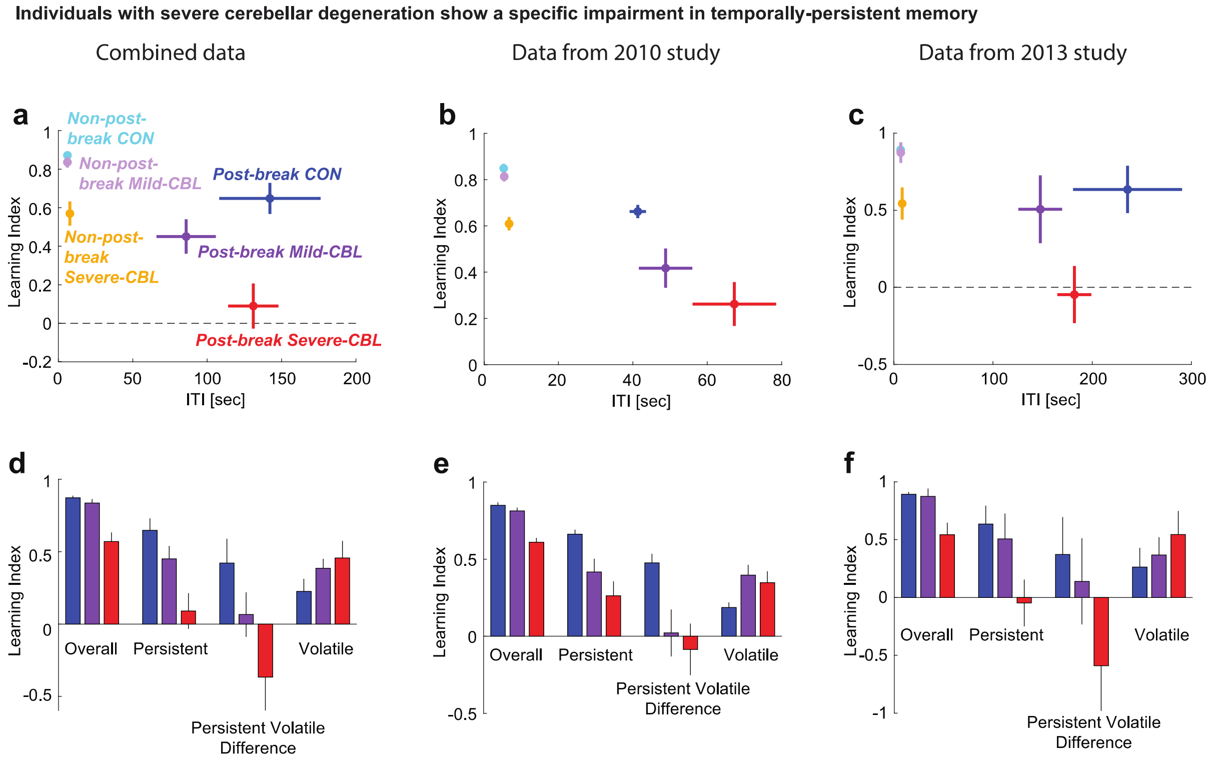


**Figure S1: Analysis of specific impairments in temporally-persistent and temporally-volatile sensorimotor memory – including patients with mild cerebellar degeneration.**

Format as in Figure 3 from the main paper, but also including patients with mild cerebellar degeneration (Mild-CBL group, ICARS<40; n=5 in the 2010 study and n=3 in the 2013 study; shown in purple).

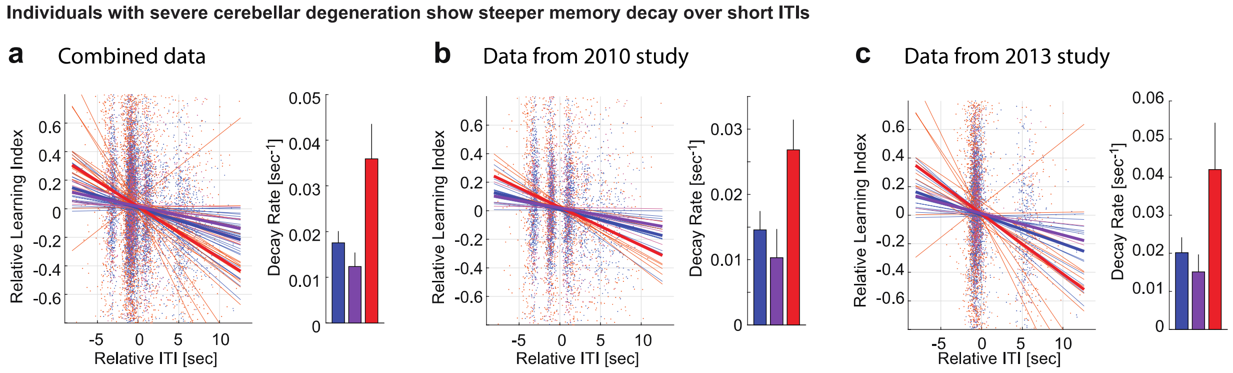


**Figure S2: Analysis of temporal decay rates for sensorimotor memory – including patients with mild cerebellar degeneration.**

Format as in Figure 4 from the main paper, but also including patients with mild cerebellar degeneration (Mild-CBL group, ICARS<40; n=5 in the 2010 study and n=3 in the 2013 study; shown in purple).
